# Assessing Mozambican Honey Quality: Differential Physical, Chemical and Microbiological Characteristics in Formal and Informal Markets

**DOI:** 10.1101/2025.08.24.672021

**Authors:** Jacob Muianga, Wilma Custódio Fumo, Samuel Bila, Emelda Simbine-Ribisse, Abílio Changule, Otilia Tomo, Manuel Garcia-Herreros, Custódio Gabriel Bila, Júlio Come

## Abstract

Honey is a natural product made by bees and is widely valued as a healthy food. Its quality can deteriorate due to microbial activity and physical or chemical changes. This study assessed honey from formal (n=12) and informal (n=12) markets in three Mozambican provinces: Maputo, Sofala, and Inhambane, to evaluate its quality and safety. Physicochemical parameters such as viscosity, water activity (Aw), pH, ash content, total soluble solids, and diastase activity were measured using standard AOAC methods. Microbiological quality was evaluated by counting aerobic mesophilic bacteria, moulds and yeasts, and by checking for pathogenic bacteria, including *Escherichia coli* and *Salmonella* species. The results showed marked differences. Honey from informal markets had a significantly higher ash content (0.613% ± 0.079%), which is a recognized indicator of contamination (p < 0.05). In contrast, formal market samples demonstrated superior quality with greater total soluble solids (77.84 °Brix ± 0.75°Brix) and viscosity (2.168 ± 0.755 Pa.s). All honey samples from formal markets retained diastase activity, while 25% (3 out of 12) of informal market samples showed no enzymatic activity. From a microbiological perspective, all samples were within safe limits. In conclusion, the physicochemical irregularities observed in honey from informal markets underline the urgent need for better hygiene practices and stronger regulatory enforcement to protect consumers and strengthen the local honey industry.

## 1. Introduction

Honey and other bee products are valued for their nutritional and therapeutic benefits and are consumed across age groups and cultural contexts^1,2^. Over time, beekeeping has shifted from extractive and destructive practices to more sustainable and rational systems, reducing environmental harm and improving bee welfare^3^.

Although honey is widely regarded as a natural and clean product, its quality is strongly influenced by factors such as floral origin, climate, and hygiene during harvesting, processing and storage^3,4^. International standards, including the *Codex Alimentarius*, recommend assessing parameters such as moisture, sugar profile (fructose, glucose, sucrose), water-insoluble solids, acidity and diastase activity, alongside microbiological analyses. Compliance with good beekeeping and hygiene practices is essential to protect public health and to ensure access to both domestic and export markets^5^.

Recent studies from Africa and Latin America have found that honey from informal supply chains is more likely to have fungal contamination, adulteration and deviations in physical and chemical properties^4,6^. These issues are often linked to poor hygiene and limited regulatory oversight^7^. A systematic review by Mesele et al.^1^ on the physical and chemical properties of honey in Eastern Africa found several cases of non-compliance. Honey from Ethiopia and Sudan exceeded accepted sucrose limits, while samples from Sudan also showed non-compliance for free acidity. In addition, honey from Uganda failed to meet accepted limits for hydroxymethylfurfural (HMF) and diastase activity set by European and Codex standards.

Mozambique is an important honey-producing country, where the sector supports rural livelihoods and informal trade^8–10^. However, its honey quality control systems remain underdeveloped. Routine laboratory testing is rare, and the country lacks national regulations on honey quality, creating a pressing need for better quality assurance.

While research on honey often focuses on its antioxidant, antimicrobial and other functional properties, there is limited information on the physical, chemical and microbiological characteristics of honey produced in Mozambique. A literature review identified only two relevant studies: Zandamela^8^, which examined honey quality in four provinces (Maputo, Inhambane, Manica and Sofala), and Serem and Bester^6^, who studied the physical, chemical and antioxidant properties of honey from southern Africa, including Maputo province. Zandamela^8^, reported lower quality in honey from small-scale beekeepers and highlighted the need for improved production conditions.

This study addresses the need to better understand the quality and safety of Mozambican honey. Its objectives were to: (i) evaluate the physical, chemical and microbiological quality of honey from formal and informal markets in three provinces; (ii) compare the findings with national and international standards; and (iii) identify risk factors for non-compliance and propose recommendations to improve quality and market competitiveness.

We anticipated that honey from informal markets would have lower quality due to weaker hygiene, poor processing methods and limited quality control. We also expected differences among provinces (Maputo, Inhambane and Sofala), reflecting variations in production practices, environmental conditions and climate.

## 2. Materials and methods

### 2.1. Experimental design and sampling strategy

This randomized cross-sectional study was conducted in 2021. A two-factor factorial experiment was conducted using a 2×3 design. The first factor (Factor A) was market type, categorized into two levels: formal markets and informal markets. The second factor (Factor B) was geographical area, comprising three levels corresponding to three distinct provinces within the study area. This factorial design facilitated the simultaneous evaluation of the main effects of market type and provinces, as well as the investigation of potential interaction effects between market type and provinces on the response variables.

The study area was selected using a Simple Random Sampling technique, specifically employing a Table of Random Numbers. The selected provinces included Maputo and Inhambane (southern Mozambique), and Sofala (central Mozambique), as shown in Fig 1. Honey collection sites (both formal and informal) were chosen prioritizing ease of access for fieldwork execution.

**Fig 1.**
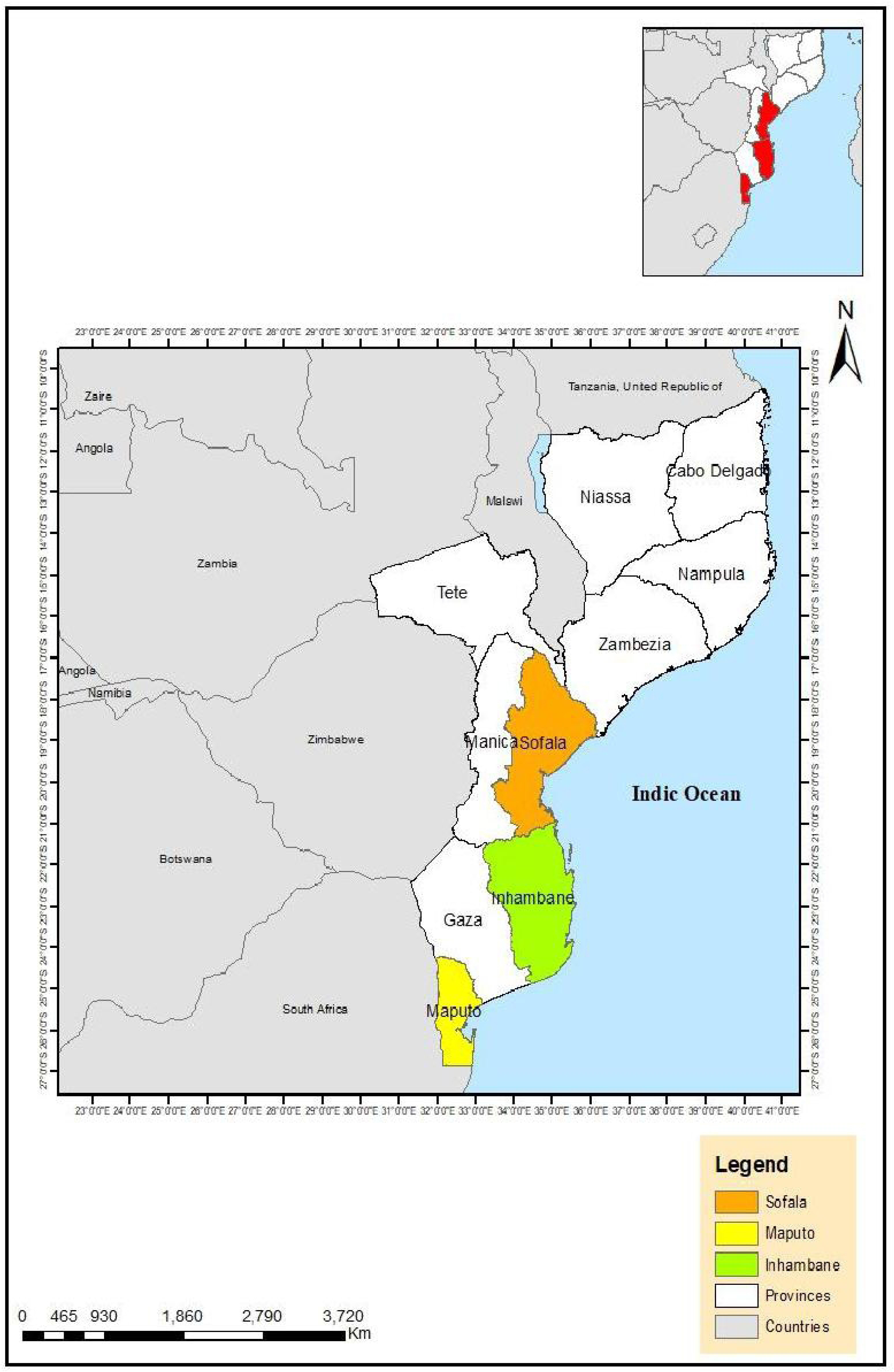
Map of Mozambique highlighting the study provinces for honey sample collection. An inset map (upper right corner) shows the location of the study provinces within the national context of Mozambique (highlighted in red). The study provinces – Sofala (orange), Maputo (yellow), and Inhambane (light green) – are highlighted. Other provinces of Mozambique are represented in white, while neighboring countries are shown in gray. The Indian Ocean is visible on the right.

### 2.2. Honey sampling

A total of 24 honey samples were purchased from three provinces in Mozambique: Maputo, Sofala and Inhambane. In Maputo, 12 samples were collected, with six from each market type. In Inhambane and Sofala, six samples were collected from each province, with three from each market type. This approach yielded 12 samples per group (n = 12), capturing variation across locations and vendors. All samples were stored in sterile glass jars at room temperature and analyzed within seven days of purchase. Aseptic techniques were used throughout collection and handling to reduce the risk of contamination.

### 2.3. Physical and Chemical Analysis

To ensure consistency and reproducibility, physical and chemical analyses for all samples were performed in duplicate using methods from the Official Methods of Analysis^11^, the International Honey Commission^12^ and the Codex Standard for Honey-CODEX STAN 12-1981^5^. The parameters measured were pH, ash content, water activity (Aw), diastase activity, total soluble solids and viscosity.

#### 2.3.1. pH

The pH of each honey sample was measured by direct potentiometric reading with a calibrated digital pH meter (Biobase, China). A 10 g aliquot of honey was used for each measurement, which was carried out at room temperature (25 °C ± 2 °C). The pH meter was calibrated before use with buffer solutions of pH 7.0 and 4.0^11^.

#### 2.3.2. Ashes

Ash content was determined by the gravimetric method described by the International Honey Commission^12^. Porcelain crucibles were first dried in a muffle furnace at 550 °C until they reached constant weight, then cooled in a desiccator. About 10 g of honey was weighed into each pre-dried crucible and incinerated in the muffle furnace at 550 °C until constant weight was reached. The crucibles were cooled in a desiccator for 20 minutes and immediately weighed. Ash content, expressed as a percentage, was calculated using equation 1:

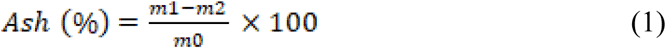

where: m0 – weight of sample honey

m1- Weight of crucible with ash

m2- weight of crucible

#### 2.3.3. Water activity (Aw)

Water activity (Aw) was measured following AOAC^11^ procedures. For each reading, 7.5 mL of honey was placed in the sample cup of a calibrated water activity meter (AquaLab-Decagon, USA). The Aw value was recorded directly.

#### 2.3.4. Total soluble solids (TSS)

Total soluble solids (TSS) were measured by direct reading with a digital refractometer (ABBE refractometer) following AOAC^11^ procedures. The instrument was calibrated with distilled water before use. About three drops of homogenized honey were placed on the prism, and the TSS value was recorded. Results were expressed in degrees Brix (°Brix).

#### 2.3.5. Viscosity

Viscosity was measured at 25 °C using a Brookfield digital viscometer (Visco 88) calibrated with distilled water. A 10 mL honey sample was read 30 seconds after spindle rotation began^13^.

#### 2.3.6. Diastase activity

Diastase activity was assessed using the iodine–starch colourimetric test^14^. A green-olive or brown colour indicated active enzymes (unheated honey), while a blue colour indicated enzyme denaturation or adulteration. Due to limited access to spectrophotometric equipment, a semi-quantitative iodine–starch method was applied, allowing classification as active or inactive rather than determining absolute diastase values in Schade units.

### 2.4. Microbiological analysis

All microbiological analyses followed the procedures of the International Commission on Microbiological Specifications for Foods^15^ and the National Laboratory of Water and Food Hygiene of Mozambique^16^.

#### 2.4.1. Quality indicator microorganisms

Quality indicator microorganisms were analyzed to assess honey quality, specifically mesophilic aerobic bacteria, moulds and yeasts. For each sample, 10 g of honey was transferred to 90 mL of peptone water and homogenized (10⁻¹). Serial decimal dilutions (10⁻²) were prepared with the same diluent. Aerobic mesophilic bacteria (AMB) were enumerated on Nutrient Agar incubated at 37 °C for 24 hours. Moulds and yeasts (MY) were counted on Potato Dextrose Agar incubated at 30 °C for 78 hours. Colonies were counted using a colony counter (Stuart, UK) and results were expressed as colony forming units per gram (CFU g⁻¹).

#### 2.4.2. Pathogenic microorganisms

Pathogenic microorganisms tested included *Escherichia coli* and *Salmonella* spp. For each, a pre-enrichment step was carried out by adding 25 g of honey to 225 mL of the specific medium.

For *E. coli*, enrichment was done in Luria Broth. Aliquots of 0.1 mL from the culture were streaked onto MacConkey agar plates using sterile plastic loops, followed by incubation at 37 °C for 24 hours in a bacteriological incubator.

For *Salmonella* spp., a standard three-stage enrichment procedure was used. Samples were pre-enriched in Lactose Broth (37 °C, 24 hours), then selectively enriched in Selenite-Cystine broth (1:10 dilution, 37 °C, 24 hours). Aliquots of 0.1 mL from the enriched cultures were streaked onto duplicate Xylose Lysine Deoxycholate (XLD) agar plates and incubated at 37 °C for 24 hours. Plates were examined for colonies characteristic of *Salmonella*, appearing pink with a black centre.

Positive and negative controls were included to validate the detection of *E. coli* and *Salmonella* spp.

### 2.5. Statistical analysis

Statistical analyses followed standard protocols for checking assumptions and selecting appropriate tests. Data normality was assessed with the Shapiro–Wilk test for each experimental group, and homogeneity of variances was evaluated with Levene’s test. Independence of observations was ensured by the randomized experimental design.

Comparisons between market types were made using independent samples t-tests when parametric assumptions were met, and Mann–Whitney U tests when they were not. Comparisons among provinces used one-way analysis of variance (ANOVA) for parametric data and Kruskal–Wallis tests for non-parametric data. The factorial design was analyzed with two-way ANOVA to assess the main effects of market type and province, as well as their interaction.

Microbiological counts were log-transformed to reduce skewness. Diastase activity, being categorical, was summarized descriptively. Statistical significance was set at p < 0.05. Analyses were performed in IBM SPSS Statistics (version 27), and descriptive summaries and visualizations were prepared in Microsoft Excel 2016.

## 3. Results

This section presents the physical, chemical and microbiological quality results for honey samples (n = 24) collected from formal and informal markets in the provinces of Maputo, Inhambane and Sofala.

### 3.1. Physical and Chemical Analysis

#### 3.1.1. Comparison of market type

Descriptive statistics showed clear differences between formal and informal markets, with ash content and viscosity displaying the largest variation between groups Table 1. Viscosity variability was notably higher in formal markets (CV = 34.8%) than in informal markets (CV = 17.0%).

**Table 1.** Descriptive statistics for formal and informal markets.

| Parameter | Market type | N | Mean ± S.E.M | Minimum | Maximum | CV (%) |
| --- | --- | --- | --- | --- | --- | --- |
| **pH** | Formal | 12 | 3.64± 0.005 | 3.62 | 3.67 | 0.40 |
| | Informal | 12 | 3.63 ± 0.006 | 3.60 | 3.65 | 0.50 |
| **Aw** | Formal | 12 | 0.62 ± 0.003 | 0.61 | 0.63 | 1.40 |
| | Informal | 12 | 0.62 ± 0.003 | 0.61 | 0.64 | 1.70 |
| **Ashes (%)** | Formal | 12 | 0.27 ± 0.005 | 0.24 | 0.29 | 5.80 |
| | Informal | 12 | 0.61 ± 0.023 | 0.49 | 0.75 | 12.8 |
| **TSS (°Brix)** | Formal | 12 | 77.84± 0.217 | 76.40 | 79.30 | 1.00 |
| | Informal | 12 | 76.42 ± 0.403 | 72.50 | 77.50 | 1.80 |
| **Viscosity (Pa.s)** | Formal | 12 | 2.17 ± 0.218 | 1.14 | 3.19 | 34.80 |
| | Informal | 12 | 1.35 ± 0.066 | 0.92 | 1.65 | 17.00 |

No statistical differences were observed between treatments (p< 0.05).

Fig 2 compares the physical and chemical characteristics of honey from formal and informal markets. Three parameters, ash content, total soluble solids (TSS) and viscosity, showed statistically significant differences between the two market types.

**Fig 2.**
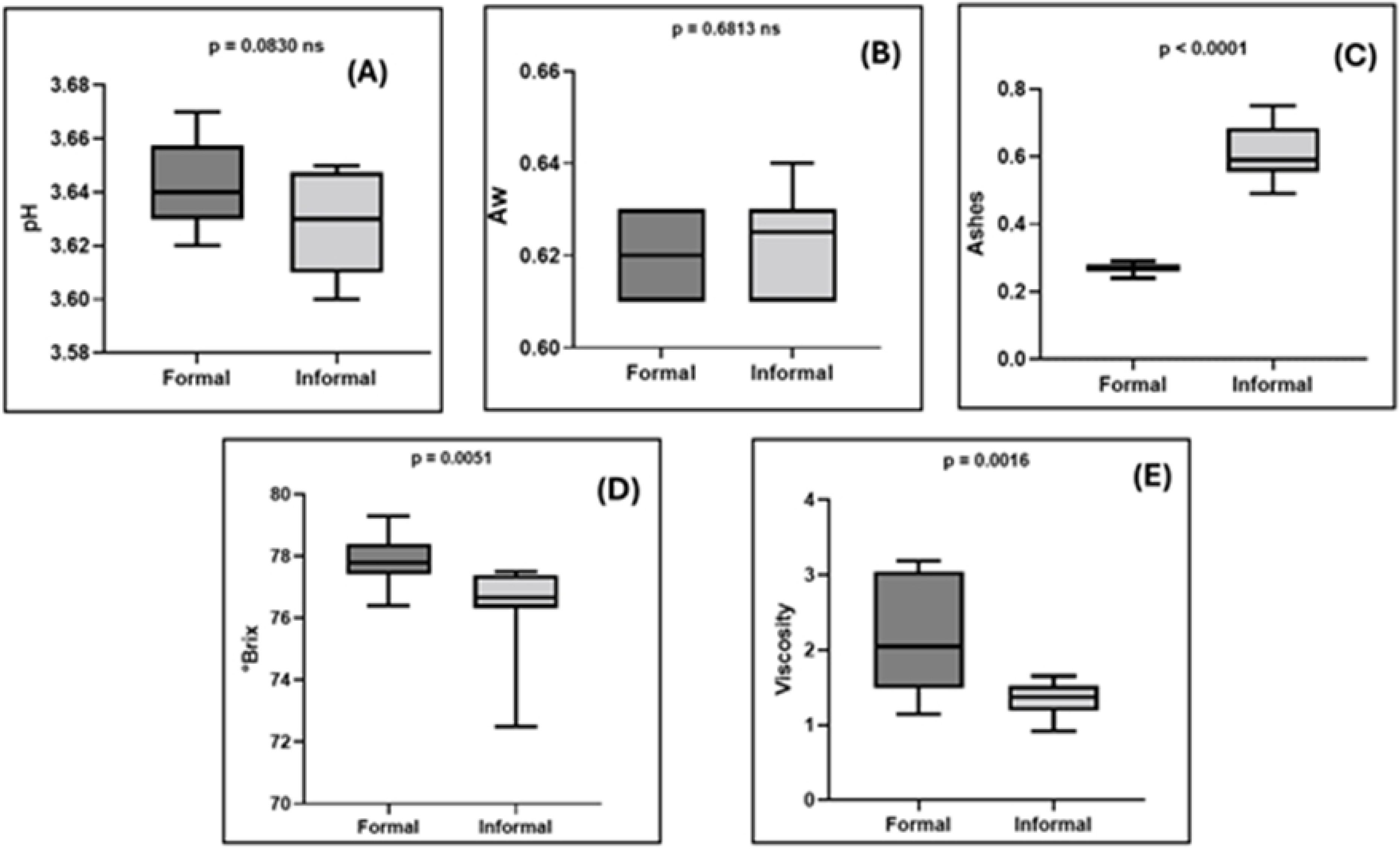
Comparative analysis of physical and chemical parameters in honey samples from formal and informal markets. (A) pH, (B) water activity (Aw), (C) ash content (%), (D) total soluble solids (°Brix) and (E) viscosity. Ns = not significant. Statistical differences were considered at p < 0.05.

Ash content was higher in informal market samples (0.613 ± 0.079%) than in formal market samples (0.269 ± 0.016%) (p < 0.001, d = –6.06). TSS was greater in formal market samples (77.84 ± 0.75 °Brix) than in informal market samples (76.42 ± 1.40 °Brix) (p < 0.05, d = 1.27). Viscosity was also higher in formal market samples (2.168 ± 0.755) compared with informal market samples (1.351 ± 0.229) (p = 0.002, d = 1.47). No significant differences were found for pH (p = 0.083) or water activity (Aw) (p = 0.681).

In the diastase enzyme test, all formal market samples (100%, 12/12) showed active diastases, indicating no enzyme denaturation. In the informal market, 75% (9/12) of samples also showed active diastase, while 25% (3/12) showed no activity.

#### 3.1.2. Comparison of analyzed parameters between geographical areas

The geographical area analysis showed moderate variability between the three provinces studied, with Maputo showing greater variability in several variables due to the larger sample size and the inclusion of both types of market.

Fig 3 illustrates the distribution of physical and chemical parameters in honey samples among the three provinces. Comparative statistical analysis demonstrated that none of the evaluated parameters exhibited significant variation among the three provinces under investigation (p > 0.05 for all comparisons).

**Fig 3.**
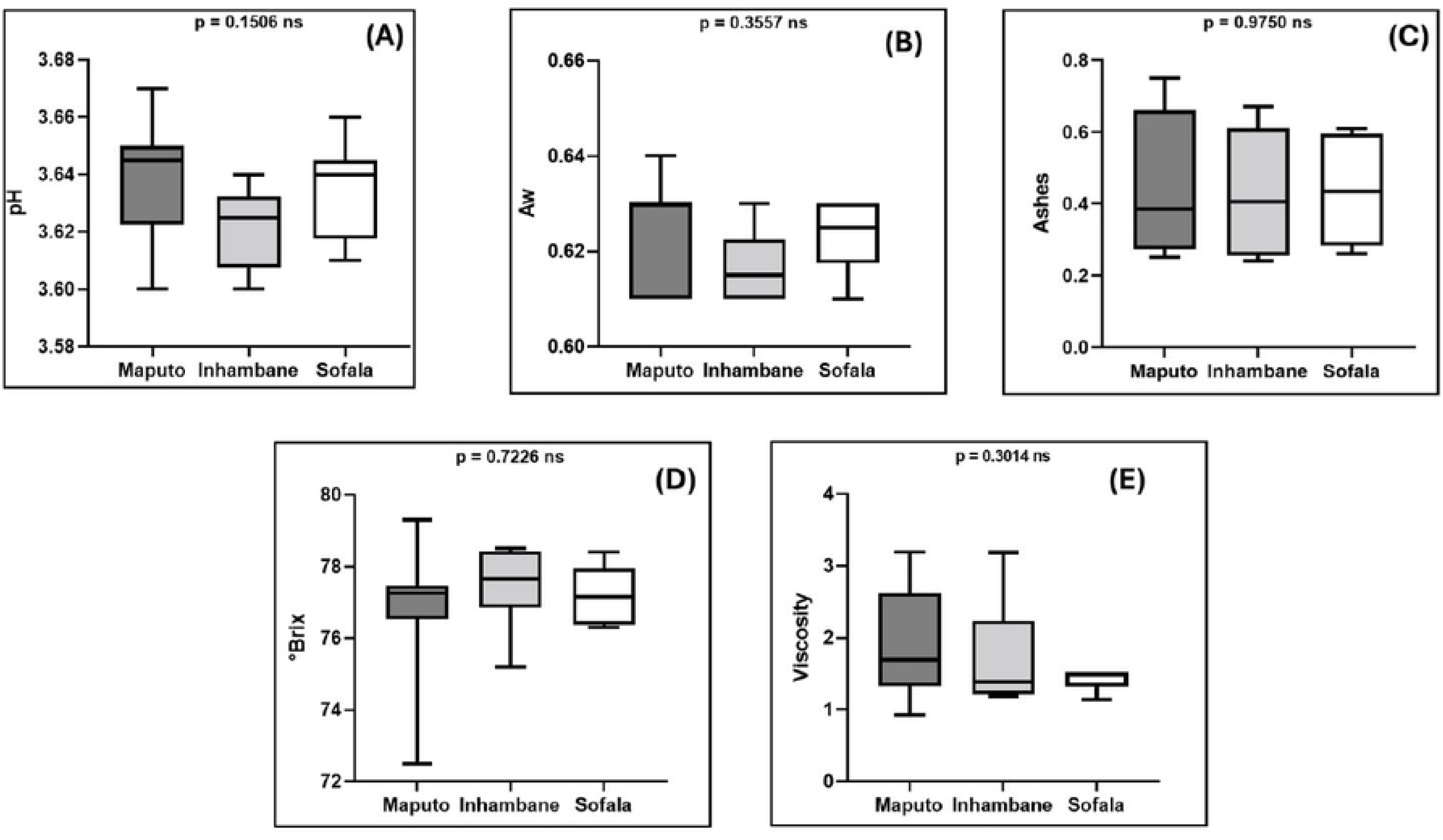
Comparative analysis of physical and chemical parameters in honey samples from three provinces of Mozambique. (A) pH, (B) water activity (Aw), (C) ash content (%), (D) total soluble solids (°Brix) and (E) viscosity. Ns = not significant. Statistical differences were considered at p < 0.05. Regarding the diastase enzyme test, among the three provinces evaluated, Maputo was found to have the most positive samples (25%, 3/12).

### 3.2. Microbiological analysis

This section presents the results of the microbiological analysis comparing the types of markets and geographical areas studied.

#### 3.2.1. Pathogenic microorganisms

Analysis of pathogenic microorganisms revealed that all honey samples showed *E. coli* and *Salmonella* spp. counts below the detection limit (<10 MPN/mL). No significant differences were observed between markets or among the three provinces.

#### 3.2.2. Quality indicators microorganisms

Fig 4 shows aerobic mesophilic bacteria and mould and yeast counts by market type and province, presented as box plots with median, interquartile range and outliers. Counts were significantly higher in formal markets for both aerobic mesophilic bacteria (1.21 ± 0.61 × 10² CFU/g; p = 0.0008) and moulds and yeasts (8.63 ± 0.44 × 10 CFU/g; p = 0.0008). No significant differences were found among provinces for aerobic mesophilic bacteria (p = 0.9336) or for moulds and yeasts (p = 0.3748).

**Fig 4.**
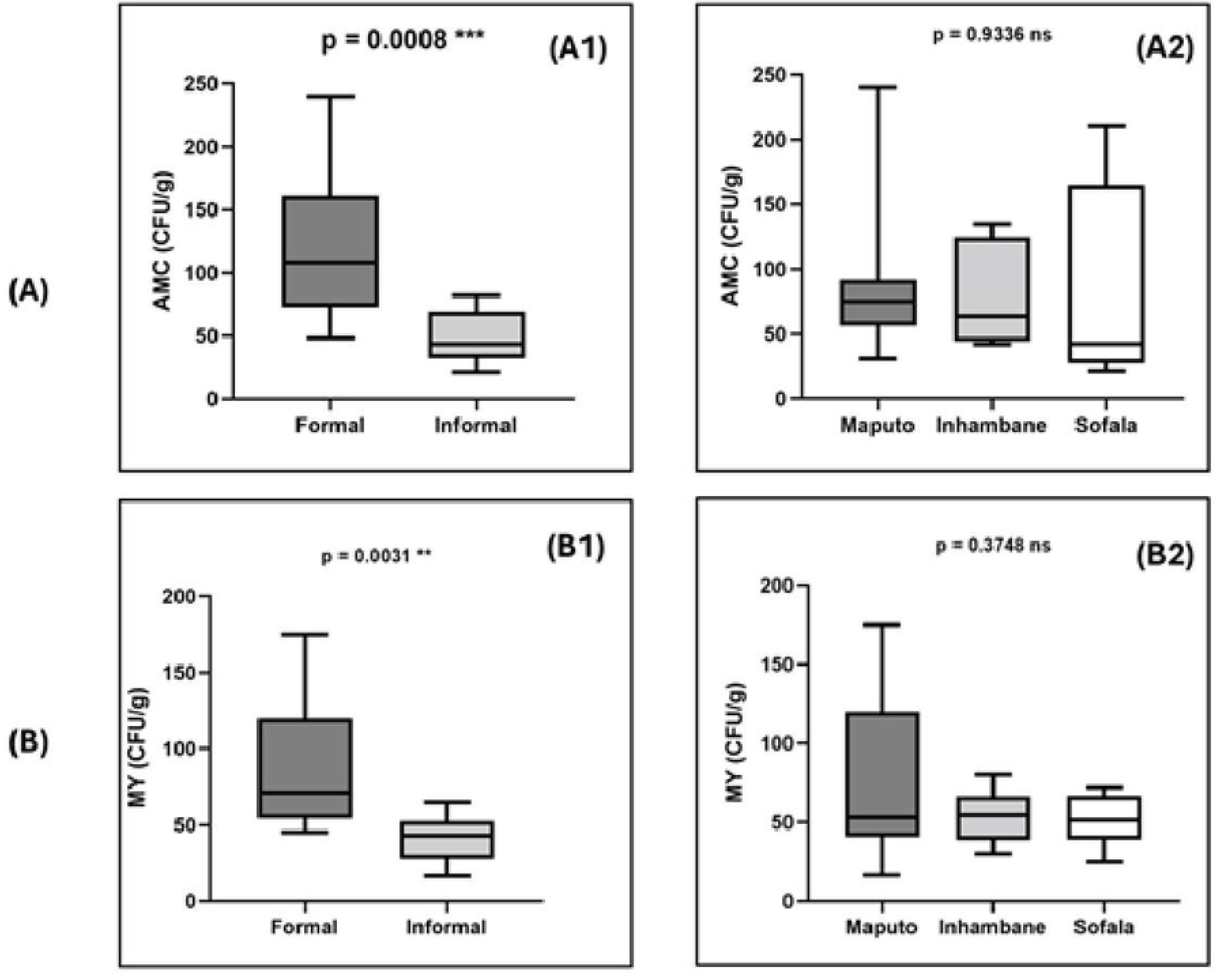
Microbiological quality parameters of honey samples from formal and informal markets in three provinces of Mozambique. (A) Aerobic mesophilic count (AMC): (A1) comparison between market types, (A2) comparison between provinces. (B) Moulds and yeasts (MY): (B1) comparison between market types, (B2) comparison between provinces. Ns = not significant; * indicates a significant difference (p < 0.005).

### 3.3. Analysis of market type × geographical area interactions

#### 3.3.1. Physical and chemical interactions

Two-way ANOVA showed a significant interaction between market type and province for viscosity (p = 0.018), with smaller differences between formal and informal markets in Sofala compared with the other provinces (Fig 5). No significant interactions were found for the other physical and chemical parameters (p > 0.05), indicating that the main effects operated independently across the geographical areas studied.

**Fig 5.**
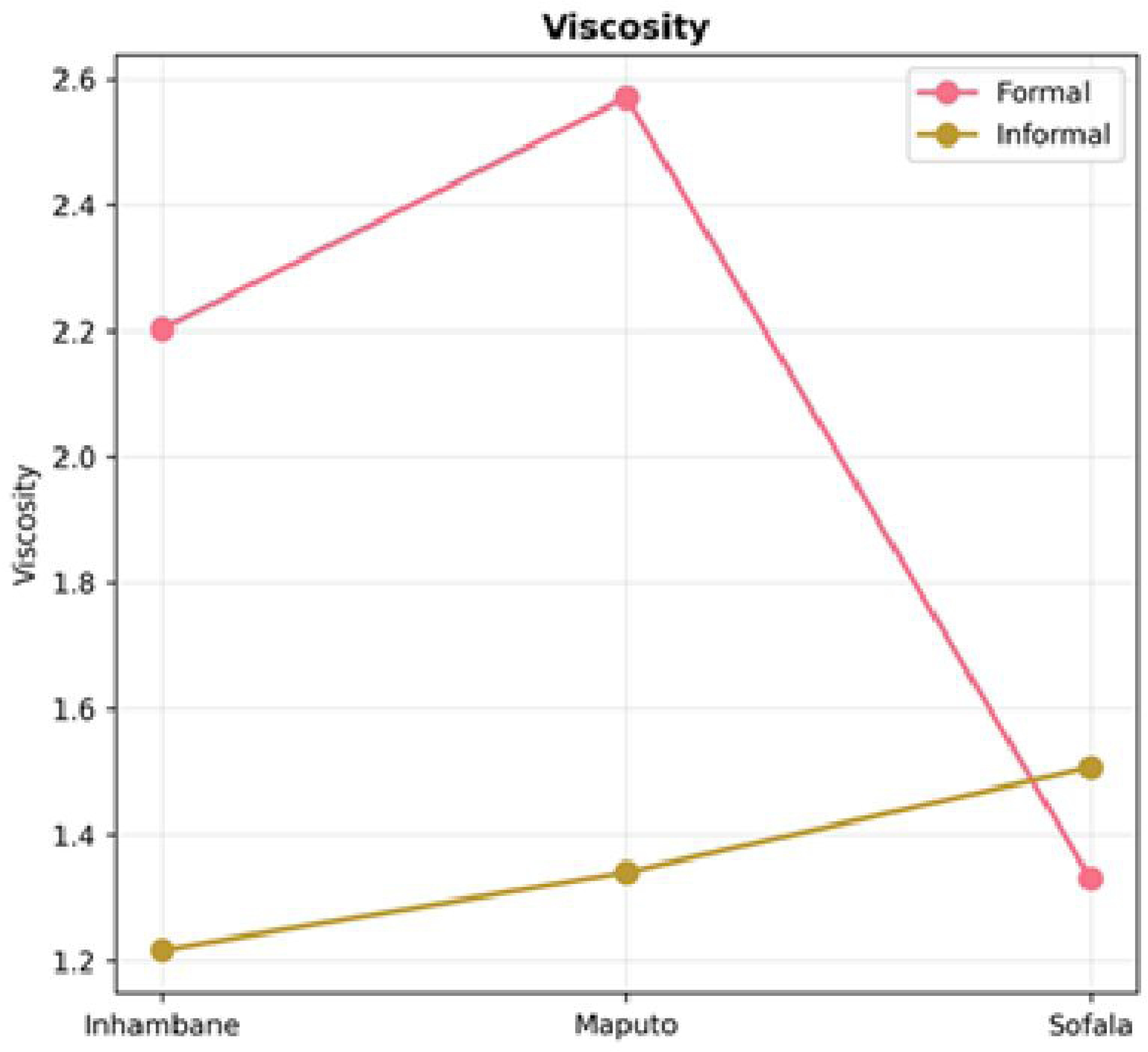
Interaction between the type of market and the study provinces for the viscosity parameter.

#### 3.3.2. Microbiological interactions

No significant interaction was observed between market type and province in the microbiological analyses for any of the parameters evaluated.

### 3.4. Correlation Coefficient Analysis

#### 3.4.1. Correlation coefficients between physical and chemical parameters

Pearson correlation coefficients were calculated for the physical and chemical parameters pH, Aw, ash content, total soluble solids (TSS) and viscosity. The correlation matrix (Fig 6) showed several statistically significant associations. Ash content was a central variable, displaying strong negative correlations with both viscosity (r = –0.66, p < 0.001) and TSS (°Brix) (r = –0.65, p < 0.001). A moderate positive correlation was also found between TSS and viscosity (r = 0.50, p < 0.05).

**Fig 6.**
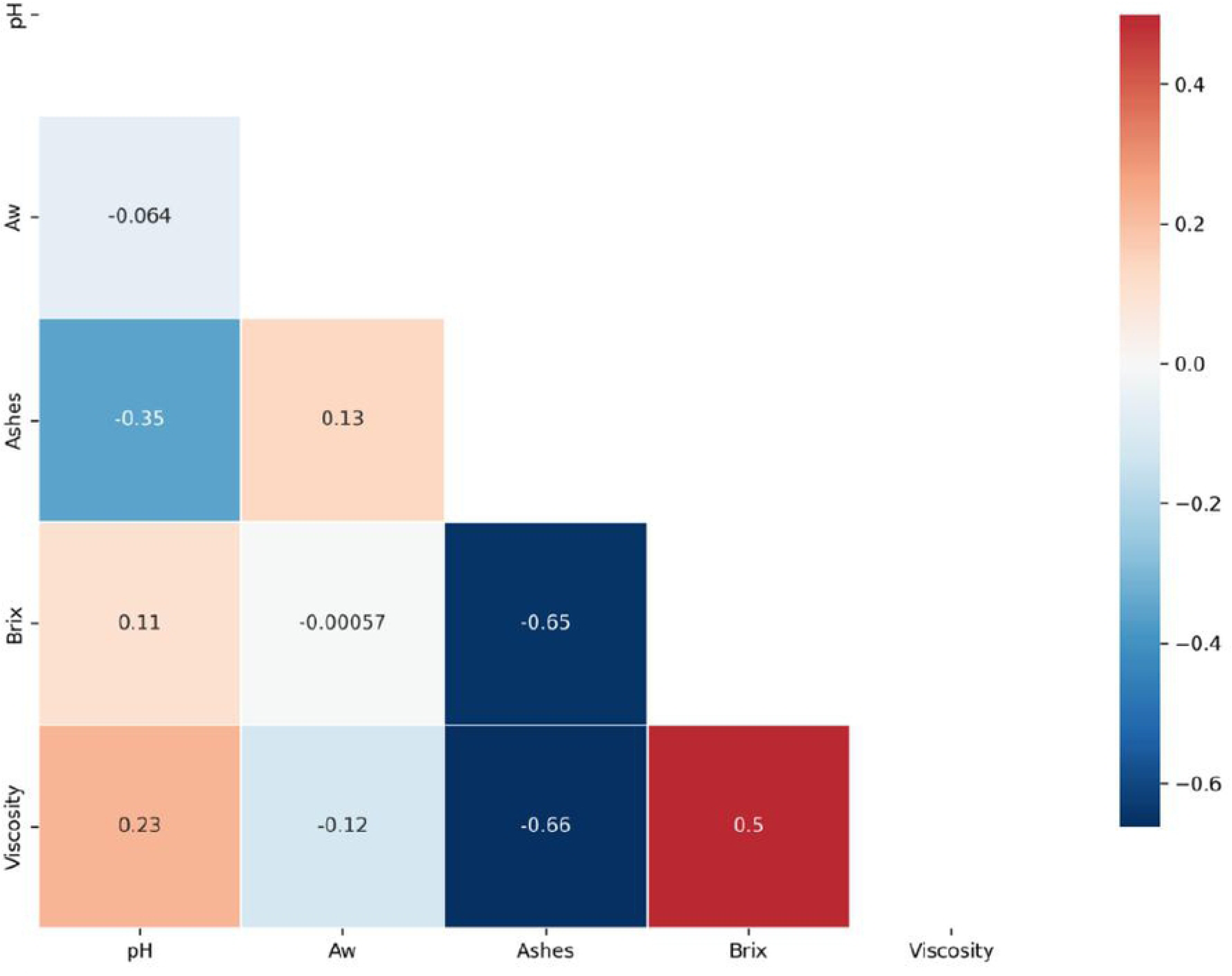
Correlation matrix heatmap of physical and chemical parameters. The colour scale represents correlation strength: dark blue = strong negative correlation, light blue = moderate negative correlation, white/grey = weak correlation, light red = moderate positive correlation and dark red = strong positive correlation.

#### 3.4.2. Correlation coefficients between microbiological analyses

Pearson’s coefficient showed a strong positive correlation between the quality indicator microorganisms AMB and MY (r = 0.566, p = 0.0040). The correlation matrix is shown in Fig 7.

**Fig 7.**
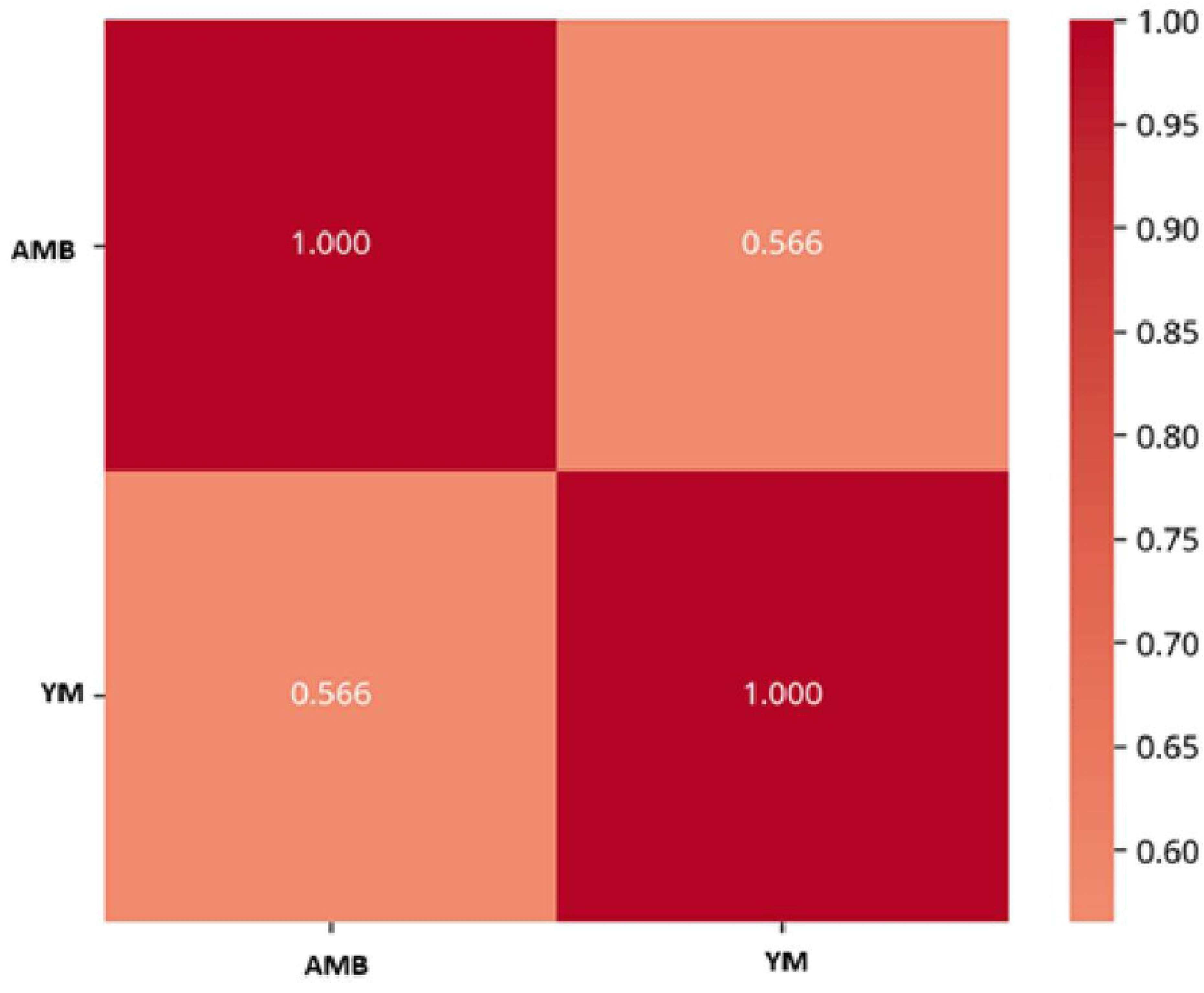
Correlation matrix of AMB and YM. Colour intensity corresponds to the correlation coefficient, as shown in the colour bar on the right. Darker red shades indicate stronger positive correlations, while lighter shades indicate weaker positive correlations.

## 4. Discussion

This section examines the physical, chemical and microbiological characteristics of honey collected from formal and informal markets in three provinces of Mozambique. The initial hypothesis predicted that honey sold through informal channels would show more non-compliance with international quality standards, mainly due to weaker hygiene and processing practices. The results partly support this prediction. While honey from both market types generally met acceptable standards, informal market samples showed higher microbial loads, greater variation in ash content, °Brix and viscosity, and some evidence of enzyme deterioration. These findings point to the need for improved handling practices and stronger quality assurance within informal value chains.

The second hypothesis, that honey quality would differ among provinces, was not supported. No significant differences were found for any parameter across regions. This result contrasts with expectations based on differences in regional infrastructure and development, suggesting that market type has a greater influence on honey quality than geographical location. Each parameter is discussed in detail below, in relation to relevant standards and the scientific literature.

### 4.1. Physical and Chemical Analyses

Although pH is not a regulated parameter, it is an important quality indicator (Pereira et al., 2020). Lower pH values can help inhibit microbial activity and extend shelf life ^1^ and may provide clues about fermentation risk or adulteration^17^. In this study, all honey samples were acidic and in line with international standards^18^. This indicates appropriate acidity for both market types. Honey acidity is influenced by floral source, nectar composition and soil characteristics^19^. Our results match established patterns reported in the literature: Serem and Bester^6^ recorded 3.87–5.12 for southern African honeys, Zandamela^8^ reported 4.11–4.42, Massingue Júnior^20^ reported 3.28–4.30, and Tanleque-Alberto et al.^9^ found 3.5–4.5 in Mozambican honeys. Comparable ranges have been reported elsewhere, including Ethiopia, with 3.55–4.05^21^, Portugal with 3.2–4.4^22^ and Brazil with 2.7-4.4^23^.

Water activity (Aw) was consistent across all samples, indicating effective moisture control. Values were similar for Maputo and Inhambane, while Sofala showed slightly higher averages, although statistical tests revealed no significant differences between market types or provinces. All values fell within the high-quality range (0.54–0.75) defined by Gomes et al.^24^. These results agree with previous Mozambican findings: Zandamela^8^ reported 0.56–0.60, and Tanleque-Alberto et al.^9^ reported 0.55–0.69. Some studies have found higher Aw values, such as Mokaya et al.^25^ with 0.70–0.77. Although Aw is not mandated in international honey standards, it is a key indicator of microbial stability. The low Aw values observed here support honey’s natural resistance to spoilage and pathogenic bacteria, contributing to its long shelf life^17^.

Total soluble solids (°Brix) differed between market types, with higher values in formal markets, but no significant differences among provinces. This suggests that geographical origin did not influence sugar concentration. Our findings are generally in line with the literature, though values vary across regions. Zandamela^8^ reported higher averages in Mozambique (82.47 °Brix in Sofala, 81.45 in Inhambane, 81.38 in Maputo), possibly due to methodological or seasonal factors. International studies show a broader range: Sant’Ana et al.^26^ found 70.0–77.7 °Brix in Brazilian semi-arid honeys, and Mokaya et al.^25^ reported 64.1–76.5 °Brix in Afrotropical bee honey. Although °Brix is not a regulated parameter, it helps estimate sugar content and detect adulteration. The higher °Brix in formal markets may reflect better concentration control or less dilution, while the uniformity across provinces suggests a consistent composition regardless of location^27^.

Viscosity showed one of the most marked differences between market types, being about 60% higher in formal markets. While not a regulated parameter, viscosity influences consumer perception of texture and can indicate authenticity and freshness^27,28^. The higher viscosity in formal market samples may be linked to lower water activity and higher sugar content, as well as better moisture control during processing and storage. These values are comparable to those found SANTOS et al.^29^ reported in the Brazilian honey viscosity (1.90 to 8.55 Pa.s).

Ash content, a key quality and purity indicator influenced by floral origin, processing practices and environmental contamination^30^, showed the largest difference between market types. Informal market samples had ash levels 128% higher than formal market samples, with averages exceeding both the Mozambican national standard (0.35%, NM 20/2005) and the *Codex Alimentarius* limit of 0.6%^5^. Elevated ash suggests contamination with dust, sand, pollen debris, or insect residues, likely due to inadequate filtration and poor handling during harvesting, processing, and storage. These findings agree with Zandamela^8^, who linked high ash content to non-compliance with Good Hygiene Practices. Half of the informal market samples exceeded international standards, raising concerns about consumer safety and limiting export potential.

Contrary to earlier Mozambican data from Zandamela^8^, which showed significant regional differences (Sofala highest at 0.63%, Maputo lowest at 0.34%), our study found no variation among provinces. This suggests that market type has a stronger influence on ash content than geographical origin, indicating that contamination is driven more by processing and handling than by environmental or floral factors. The shift from historical patterns may reflect changes in processing methods, environmental conditions or market dynamics over the past decade.

Diastase activity results showed all formal market samples had active enzymes, while 25% (3/12) of informal market samples lacked activity. Pereira et al.^27^ observed enzyme loss in only 4.2% of samples (2/48), attributing this mainly to high pasteurization temperatures. Overheating during processing, poor storage and adulteration with inverted sugar syrup (which contains no diastases) are other possible causes^1,31^. Low diastase activity is often associated with high hydroxymethylfurfural (HMF) levels, and *Codex Alimentarius* recommends combining diastase testing with other chemical tests to confirm adulteration or detect excessive heating ^5^.

### 4.2. Microbiological analysis

A critical gap exists in Mozambique’s food safety legislation regarding microbial contamination limits and hygiene standards specific to honey. In the absence of national regulations, the country relies on international frameworks, such as *Codex Alimentarius* guidelines, or on standards from other countries. For this study, the reference limits were based on Brazilian regulations from the Ministry of Agriculture for products of animal origin^32^.

No pathogenic microorganisms were detected in any of the samples. *Escherichia coli* was absent from all honey tested, in agreement with findings from Zandamela^8^ in Mozambique and from Machado et al.^22^ and Estevinho et al.^33^ in Portugal. The absence of *E. coli* is recognized as an indicator of good sanitary quality^22^. Likewise, no colonies suggestive of *Salmonella* spp. were found, consistent with Pereira et al.^34^ and Fernández et al.^35^. The absence of pathogenic bacteria in our study indicates that honey from both market types and all provinces met basic food safety standards. This finding is consistent with honey’s natural antimicrobial properties, which are linked to its low water activity and acidic pH.

Assessment of indicator microorganisms — aerobic mesophilic bacteria (AMB) and moulds and yeasts (MY) — revealed an unexpected pattern. Formal market samples had higher counts than informal market samples, despite their presumed advantages in hygiene practices and lower water activity. All counts, however, were within acceptable limits under Brazilian regulations^32^. The unexpected results warrant further scrutiny of analytical and sampling procedures, as they may reflect laboratory artefacts rather than true quality differences. Possible explanations include sample mix-ups, laboratory contamination or deviations from standard procedures. Quality control and blank samples should be reviewed to rule out such issues.

The microbial counts observed here are comparable to those reported by Machado et al.^22^, with aerobic mesophilic bacteria < 10³ CFU/g and yeasts and moulds < 10² CFU/g, both within acceptable criteria. Zandamela^8^ likewise found AMB and MY values within legal limits.

AMB counts are widely used as indicators of hygiene, reflecting conditions during harvesting, handling, processing and storage. These results may be influenced by intrinsic factors such as pH and water activity, and by extrinsic factors such as temperature and environmental contamination throughout production and distribution While the overall results meet safety requirements, they do not confirm full compliance with good hygiene or processing practices. The detection of excessive mould and yeast counts in some informal market samples points to the need for improved training and oversight of honey producers and vendors. Future studies should increase sample size and broaden the range of microbiological and physical and chemical parameters assessed, to build a more complete picture of honey quality in Mozambique^34,35^.

### 4.3. Interaction Market Type × geographical area

Viscosity showed a significant interaction between market type and province. In Sofala, the difference between formal and informal markets was smaller than in Maputo and Inhambane. This suggests that the impact of market type on viscosity is influenced by geographical factors, possibly reflecting regional differences in processing or storage practices that affect honey flow properties. Post-hoc analysis confirmed that the formal– informal disparity in viscosity was attenuated in Sofala compared with the other provinces.

No significant interactions were found for pH, water activity, ash content or total soluble solids, indicating that market type and province act independently for these parameters. This consistency supports the robustness of the main effects and suggests that market type differences are stable across geographical areas for most quality indicators.

For microbiological parameters, no interaction was detected between market type and province, indicating that variations in microbial counts are related to market type rather than geographical origin.

Previous studies on Mozambican honey quality have focused mainly on geographical variation. Zandamela^8^ reported differences between southern and central regions, while Tanleque-Alberto et al.^9^ assessed central and northern areas. However, these investigations used single factor designs that could not separate the effects of geography from those of the market system.

This study introduces a two-factor experimental approach that evaluates both geographical and market type effects simultaneously. The factorial design (2 × 3) revealed that market type is the dominant factor influencing honey quality, outweighing the geographical effects documented in earlier research. These findings challenge the prevailing view that quality differences are primarily geography-driven, showing instead that commercialization channels have a stronger impact. This shift in understanding has important implications for quality control and market development strategies in Mozambique.

### 4.4. Correlation coefficients analysis

The correlation matrix showed several statistically significant relationships among the physical and chemical parameters. Ash content was a central variable, with strong negative correlations to viscosity (r = –0.66) and total soluble solids (°Brix) (r = –0.65). A moderate positive correlation was found between °Brix and viscosity (r = 0.50).

These patterns suggest that higher ash content — potentially indicating contamination or excess mineral impurities — is linked to reduced viscosity and lower soluble solids, both of which may reflect deteriorated product quality. In contrast, the positive correlation between °Brix and viscosity is consistent with the expected technological relationship in which higher soluble solids increase product viscosity^27^.

A strong positive association was also observed between mesophilic bacteria and moulds and yeasts, possibly due to shared intrinsic factors that promote their growth, such as pH and water activity.

## 5. Conclusion

Our findings show that honey quality in Mozambique, as assessed by physical, chemical and microbiological parameters, is not influenced by geographical origin. In contrast, market type is a determining factor. Honey from formal markets generally displayed higher quality than that from informal markets and complied with international standards, particularly those of the *Codex Alimentarius*, confirming its safety for human consumption.

Honey from informal markets, while microbiologically acceptable in most cases, presented isolated non-compliances such as elevated fungal contamination and reduced diastase activity. These issues may be linked to inadequate handling practices or thermal degradation.

Viscosity was the only parameter that showed an interaction between market type and province, with differences between formal and informal markets varying according to geographical location.

## Acknowledgments

We gratefully acknowledge Intermed Mozambique Lda for their financial support in covering the publication fees for this article, enabling open access dissemination of our research findings.

